# Downregulation of miR-17-92 cluster by PERK fine-tunes unfolded protein response mediated apoptosis

**DOI:** 10.1101/2020.10.26.354894

**Authors:** Danielle E. Read, Ananya Gupta, Karen Cawley, Laura Fontana, Patrizia Agostinis, Afshin Samali, Sanjeev Gupta

**Author notes:** Corresponding author. E-mail address (S. Gupta). Equal contribution (Danielle E. Read, Ananya Gupta).

## Abstract

An important event in the unfolded protein response (UPR) is the activation of the endoplasmic reticulum kinase PERK (EIF2AK3). The PERK signalling branch first mediates a prosurvival response, which switches into a proapoptotic response upon prolonged ER stress. However, the molecular mechanisms of PERK-mediated cell death are not well understood. Here we show that expression of the primary miR-17-92 transcript and mature miRNAs belonging to miR-17-92 cluster is decreased during UPR. We found that activity of miR-17-92 promoter reporter was reduced during UPR in a PERK-dependent manner. We show that activity of miR-17-92 promoter is repressed by ectopic expression of ATF4 and NRF2. The promoter deletion analysis and ChIP assays mapped the region responding to UPR-mediated repression to site in the proximal region of the miR-17-92 promoter. Hypericin-mediated photo-oxidative ER damage reduced the expression of miRNAs belonging to miR-17-92 cluster in wild-type but not in PERK-deficient cells. Importantly, ER stress-induced apoptosis was inhibited upon miR-17-92 overexpression in SH-SY5Y and H9c2 cells. Our results reveal a novel function for NRF2, where repression of miR-17-92 cluster by NRF2 plays an important role in ER stress-mediated apoptosis. The data presented here provides mechanistic details how sustained PERK signalling via NRF2 mediated repression of miR-17-92 cluster can potentiate cell death.

## Introduction

Physiological and pathological conditions that interfere with the homeostasis of the endoplasmic reticulum (ER) can lead to the accumulation of misfolded proteins within the ER. Such an increase of misfolded proteins is known as ER stress. The unfolded protein response (UPR) constitutes a signal transduction pathway that responds to the accumulation of misfolded proteins in the ER. The UPR orchestrates an increase in ER-folding capacity through transcriptional induction of ER folding, lipid biosynthesis, and ERAD machinery along with a concomitant decrease in the rate of protein synthesis through selective mRNA degradation and translational repression [1]. The UPR is therefore cytoprotective, allowing cells to adapt to perturbations that impinge on ER protein folding. However, during severe and prolonged ER stress, the UPR can become cytotoxic and induce apoptosis [2].

In metazoans ER stress is detected by three ER resident proteins: activating transcription factor 6 (ATF6), protein kinase RNA-like ER kinase (PERK) and inositol-requiring enzyme 1 (IRE1) [3]. PERK-mediated phosphorylation of eukaryotic translation initiation factor 2α on the alpha subunit (eIF2α at Ser51 leads to translational attenuation [4]. Whilst phosphorylation of eIF2α inhibits general translation initiation, it paradoxically increases translation of activating transcription factor 4 (ATF4), which induces the transcription of genes involved in restoration of ER homeostasis [4]. The endoribonuclease activity of IRE1 is responsible for the nonconventional splicing of transcription factor X-box binding protein (XBP1), which controls the transcription of chaperones and genes involved in ER-associated protein degradation (ERAD) [5]. In response to ER stress, ATF6 traffics to the Golgi complex where S1P and S2P proteases cleave the cytosolic and transmembrane domains. The processed form of ATF6 (N-terminal transcriptional regulatory domain) translocates to the nucleus thereby regulating the transcription of genes involved in ER homeostasis, such as ER chaperones and ERAD components [1].

Although we understand the activation and general function of UPR components the molecular mechanisms of ER stress-induced cell death are not well understood. MicroRNAs (miRNAs) are a family of short (20–23 nucleotide), endogenous, single-stranded RNA molecules that regulate gene expression in a sequence-specific manner. MiRNAs are generated from primary RNA transcripts that are processed by microprocessor complex in the nucleus to generate small hairpin RNA (Precursor miRNA) which are exported to the cytoplasm. In the cytoplasm precursor-miRNA molecules undergo Dicer-mediated processing thus generating mature miRNA [6, 7]. The mature miRNA assembles into the ribonucleoprotein silencing complexes (RISCs) and guides the silencing complex to specific mRNA molecules [8]. The main function of miRNAs is to direct posttranscriptional regulation of gene expression, typically by binding to the 3’ UTR of cognate mRNAs and inhibiting their translation and/or stability [9]. MiRNAs have been implicated in nearly all developmental and pathological processes in animals such as tissue morphogenesis, cell proliferation, apoptosis, and major signalling pathways [6, 7, 10]. Global approaches in several cellular contexts have revealed that UPR regulates the expression of many miRNAs that play an important role in the regulation of life and death decisions during UPR [11, 12].

The miR-17-92 cluster comprises a group of six miRNAs on chromosome 13 that is transcribed as a single polycistronic unit [13]. Amplification and overexpression of the miR-17-92 cluster has been documented in B-cell lymphomas, lung cancer and gastric cancer [13]. The transcription factors c-MYC and E2F3 induce the expression of the miR-17-92 cluster [14] where as p53 and AML1 down regulate its expression [15, 16]. Enforced expression of miR-17-92 cluster miRNAs in an Eμ-myc transgenic mouse model of B cell lymphoma accelerates disease onset and progression [13, 17].

Deletion of the miR-17-92 cluster results in smaller embryos and immediate postnatal death of all animals, and is associated with severe lung hypoplasia and defective ventricular septum [18]. However, in contrast to the wealth of information about the biological effects of the miR-17-92 cluster, little is known about its regulation.

In this study we describe the role of the miR-17-92 cluster in ER stress responses. Our data indicate that expression of miR-17-92 cluster is reduced during conditions of ER stress in a variety of cell types. Ectopic expression of ATF4 or NRF2 leads to reduced expression of miRNAs belonging to the miR-17-92 cluster. We show that the miR-17-92 cluster is repressed by the PERK-dependent transcription factors ATF4 and NRF2. We provide evidence that repression of the miR-17-92 cluster contributes to the ER stress-mediated apoptosis. Taken together our results suggest a role for the miR-17-92 cluster during the ER stress response.

## Materials and Methods

### Cell culture and treatments

MEFs, H9c2, MDA-MB231, 293T, PC12, SH-SY5Y and MCF-7 cells were maintained in Dulbecco’s modified medium (DMEM) supplemented with 10% FCS, 100 U/ml penicillin and 100 mg/ml streptomycin at 37 °C with 5% CO2. PC12 cells were cultured in DMEM supplemented with 10% heat inactivated horse serum, 5% foetal bovine serum and 1% penicillin/streptomycin (Sigma) at 37 °C with 5% CO2. Cells were treated with thapsigargin and tunicamycin for the times indicated. All reagents were from Sigma-Aldrich unless otherwise stated.

### Plasmid constructs

The miR-17-92 cluster promoter reporter constructs (Prom17M [miR17-92FL]), Prom17 [17-92/1] and Prom17/1 [17–92] were from Dr Laura Fontana [16, 20]. The expression plasmids for ATF4, wild type PERK and K618A PERK were obtained from Dr David Ron. The expression vector for wild type NRF2 was from Dr. Alan Diehl, University of Pennsylvania, USA [23]. The expression vector for wild-type CHOP was from Dr. Andreas Strasser, WEHI, Australia [37]. For lentiviral expression of miRs-17-92, a 1-kb fragment spanning the miR17-92 cluster was amplified using pCXN2-miR17-92 plasmid [38] as a template and cloned in to pCDH-CMV-EF1-RFP. Transient transfections were carried out using Lipofectamine 2000 (Invitrogen) according to the manufacturer’s protocol.

### Generation of stable cell lines

We generated stable subclones of H9c2 and SH-SY5Y expressing miR-17-92 cluster by transducing the cells with pCDH-CMV-miR-17-92-EF1-RFP or the corresponding control pCDH-CMV-EF1-RFP. Cells were transduced with the lentivirus using polybrene (5 μg/mL) to increase the transduction efficiency. The subclones expressing miR-106b-25 cluster were obtained by sorting on the basis of RFP using the FACS AriaII cell sorter (BD) to attain >90% RFP positivity in the selected population

### RNA extraction, RT-PCR and real time RT-PCR

Total RNA was isolated using RNeasy kit (Qiagen) according to the manufacturer’s instructions. Reverse transcription (RT) was carried out with 2 μg RNA and Oligo dT (Invitrogen) using 20 U Superscript II Reverse Transcriptase (Invitrogen). Real-time PCR method to determine the induction of UPR target genes has been described previously. Briefly, cDNA products were mixed with 2 × TaqMan master mixes and 20 × TaqMan Gene Expression Assays (Applied Biosystems) and subjected to 40 cycles of PCR in StepOnePlus instrument (Applied Biosystems). Relative expression was evaluated using the ΔΔCT method.

### Measurement of miRNA Levels Using TaqMan qRT-PCR Assays

Total RNA was reverse transcribed using the TaqMan miRNA Reverse Transcription Kit and miRNA-specific stem-loop primers (Applied BioSystems) in a small-scale RT reaction [comprised of 0.19 μL of H2O, 1.5 μL of 10X Reverse-Transcription Buffer, 0.15 μL of 100 mM dNTPs, 1.0 μL of Multiscribe Reverse-Transcriptase (50 Units/μL), and 5.0 μL of input RNA (20 ng/μL); components other than the input RNA were prepared as a larger volume master mix], using a Tetrad2 Peltier Thermal Cycler (BioRad) at 16°C for 30 min, 42°C for 30 min and 85°C for 5 min. For miR-17, miR-18a, miR-19a, miR-19b, miR-20a, miR-92a snoRNA and U6 snRNA, 4.0 μL of RT product was combined with 16.0 μL of PCR assay reagents (comprised of 5.0 μL of H2O, 10.0 μL of TaqMan 2X Universal PCR Master Mix, No AmpErase UNG and 1.0 μL of TaqMan miRNA Assay) to generate a PCR of 20.0 μL of total volume. Real-time PCR was carried out using an Applied BioSystems 7900HT thermocycler at 95°C for 10 min, followed by 40 cycles of 95°C for 15 s and 60°C for 1 min. Data were analysed with SDS Relative Quantification Software version 2.2.2 (Applied BioSystems.), with the automatic Ct setting for assigning baseline and threshold for Ct determination.

### Luciferase reporter Assays

In promoter assays, MCF-7 cells were transfected with 0.8 μg of firefly luciferase vectors (empty pGL4 or pGL4prom17M vector), in combination with a *Renilla* luciferase vector (0.2 μg) as internal control. Twenty four hours post-transfection cells were treated with thapsigargin or tunicamycin for 24 h. Firefly luciferase and Renilla luciferase activities were measured 48 h after transfection at 560 nm using a 10s luminescence protocol with a Wallac plate reader and then normalized for *Renilla* luciferase activity.

### Chromatin Immunoprecipitation Assay

MCF-7 cells (3 × 106 cells) were transfected with wild type ATF4, NRF2 and GFP expressing plasmids. 24 hour post transfection chromatin immunoprecipitation assay was performed with a commercial kit (ChIP-IT enzymatic kit, Active Motif_Cat# 53006) according to the manufacturer’s protocol. In brief, cells were treated with 1% (v/v) formaldehyde at room temperature for 10 min and then quenched with glycine at room temperature. The medium was removed, and cells were harvested in lysis buffer. Following 30 mins incubation on ice, samples were passed through a 23 gauge syringe up and down along the side of tube ~30 times to release nuclei and then centrifuged at 5000 rpm for 10 min at 4°C to pellet nuclei. The nuclei pellet was resuspended in 1 ml digestion buffer with 5 μL PMSF and 5 μL PIC and pre-warmed for 5 min at 37°C. For enzymatic shearing, 50 μL of 1:100 diluted shearing enzyme was added to pre-warmed nuclei, vortexed to mix and incubated at 37°C for 30 mins. The reaction was stopped by adding 20 μL ice cold EDTA for 10 mins on ice. The sample was centrifuged at 10,000 rpm for 10 mins at 4°C and the supernatant collected. 25 μL was kept for DNA purification and confirmation of shearing on a 1% agarose gel. The remainder was aliquoted into 250 μL volumes and stored at −80°C (each aliquot can be used for 4 chip reactions). The sheared samples were precleared with Protein Gagarose beads at 4°C overnight. A small amount of chromatin (10 μL) was kept as the input sample at −20°C until the IP samples were ready. Subsequently, immunoprecipitation was conducted with anti-ATF4 antibody (Proteintech_Cat# 60035-1), anti-Nrf2 antibodies (Epitomocs_Cat# 2178-1). Normal IgG and RNA Pol II anti-bodies were used as controls (provided within the kit). Immunocomplexes were collected the following morning using Protein G-agarose beads pelleted by centrifugation and washed with low salt buffer, high salt buffer, and Tris-EDTA buffer (25 mM Tris-HCl, 150 mM NaCl, 1 mM EDTA, pH 7.2) to remove any nonspecific binding. The immunocomplexes were eluted from the beads using 50 μL of elution buffer (1 M NaHCO3, 1% SDS). The cross-links of the protein-DNA complexes were reversed by adding 4 μL of 5M NaCl and 1 μL of RNAse A followed by an overnight incubation of the eluted products at 65°C. A total of 2 μL of proteinase K (10 μg/μL) was subsequently added to the solution, and samples were incubated at 42°C for 2 h. DNA was then purified using the spin columns provided. MiR-17-92 gene promoter sequences were amplified by PCR with miR17-92Fwd: 5’-GTGTCAATCCATTTGGGAGAG-3’ and miR17-92Rev: 5’-TGGTCACAATCTTCAGTTTTAC-3’. CHOP gene promoter sequences were amplified by CHOPFwd: 5’-GGGCCAAGAAATATGGGAGT-3’ and CHOPRev: 5’-TAGTCGGTCGTGAGCCTCTT-3’. Hemeoxygenase-1 gene promoter sequences were amplified by HO-1Fwd: 5’-GCTGCCCAAACCACTTCTGT-3’ and HO-1Rev: 5’-GCCCTTTCACCTCCCACCTA-3’. The PCR products were analysed in a 1% agarose gel.

### Apoptotic nuclei measurements

Cells were grown on the coverslips and after treatment, cells were fixed in (10% v/v) formaldehyde for 10 minutes at room temperature. After washing with PBS cells were mounted on glass slides in a mountant with DAPI (Vectashield_Cat# H-1200). Nuclei were visualized using an Olympus BX61 fluorescence microscope. Apoptotic nuclei (condensed, fragmented, intensely stained) were counted and presented as a percentage of total nuclei. At least 100 cells were counted per well, and all treatments were performed in triplicate.

### Western blotting

Cells were washed once in ice-cold PBS and lysed in whole cell lysis buffer (20 mM HEPES pH 7.5, 350 mM NaCl, 0.5 mM EDTA, 1 mM MgCl2, 0.1 mM EGTA and 1% NP-40) after stipulated time of treatments and boiled at 95o C with Laemmli’s SDS-PAGE sample buffer for 5 min. Protein concentration was determined by Bradford method. Equal amount (20 μg/lane) of protein samples were run on an SDS polyacrylamide gel. The proteins were transferred onto nitrocellulose membrane and blocked with 5% milk in PBS-0.05%Tween. The membrane was incubated with the primary antibody for cleaved caspase-3 (ISIS, Cat# 9664) or β-Actin (Sigma, Cat# A-5060) for 2 h at room temperature or overnight at 4o C. The membrane was washed 3 times with PBS-0.05% Tween and further incubated in appropriate horseradish peroxidase-conjugated secondary antibody (Pierce) for 90 min. Signals were detected using Western Lightening Plus ECL (Perkin Elmer).

### Statistical Analysis

The data is expressed as mean ± SD for three independent experiments. Differences between the treatment groups were assessed using Two-tailed paired student’s t-tests. The values with a p<0.05 were considered statistically significant.

## Results and Discussion

### Expression of the pri-miR-17-92 is down-regulated upon induction of ER stress

In preliminary experiments, the relative abundance of miRNAs comprising the Sanger miRBase database (Release 11.0) were analysed by microarray (LC sciences, Houston, TX, USA) using RNA from H9c2 cells during conditions of ER stress. We observed downregulation of all six miRNAs belonging to the miR-17-92 cluster in H9c2 cells treated with thapsigargin (TG) or tunicamycin (TM). Thapsigargin, [an inhibitor of the sacroplasmic/endoplasmic reticulum Ca2+-ATPase (SERCA) pump] and tunicamycin, [an inhibitor of N-linked glycosylation] both lead to accumulation of misfolded proteins in the ER and initiate UPR [19]. Since the six miRNAs belonging to the miR-17-92 cluster are derived from the primary transcript of the miR-17-92 gene, we reasoned that the miR-17-92 gene is transcriptionally regulated during ER stress. We evaluated the expression of primary miR-17-92 transcript during conditions of ER stress. We observed that upon treatment with thapsigargin (TG) or tunicamycin (TM) the level of GRP78 increased in a time dependent manner. However, under similar conditions the levels of primary miR-17-92 transcript were reduced (Figure 1A). Glucose deprivation is one of the crucial physiologic conditions leading to UPR activation, which is associated with several human diseases including tissue ischemia and cancer [4]. H9c2 cells were subjected to a combination of serum and glucose deprivation as described in materials and methods. We observed that glucose deprivation induced the expression of UPR target genes GRP78 and HERP (UPR target genes), thereby confirming the induction of UPR (Figure 1B) upon glucose deprivation. We found that conditions of glucose deprivation decreased the levels of primary miR-17-92 transcript in H9c2 cells (Figure 1B). Further treatment of MDA-MB231 cells (Figure 1C) and HEK-293T (Figure 1D) cells with thapsigargin (TG) or tunicamycin (TM) led to induction of UPR-target genes (GRP78 and HERP) and downregulation of the primary miR-17-92 transcript. Furthermore, TM treatment of H9c2 cells showed a decrease in the expression of primary miR-17-92 transcript as well as all the six miRNAs belonging to the miR-17-92 cluster (Figure 2A-B). We have used a variety of cell lines and inducers of UPR in this study and observed reduced expression of primary miR-17-92 transcript during UPR. Our results suggest that downregulation of the miR-17-92 gene in response to ER stress is not cell or stimulus dependent.

**Figure 1.**
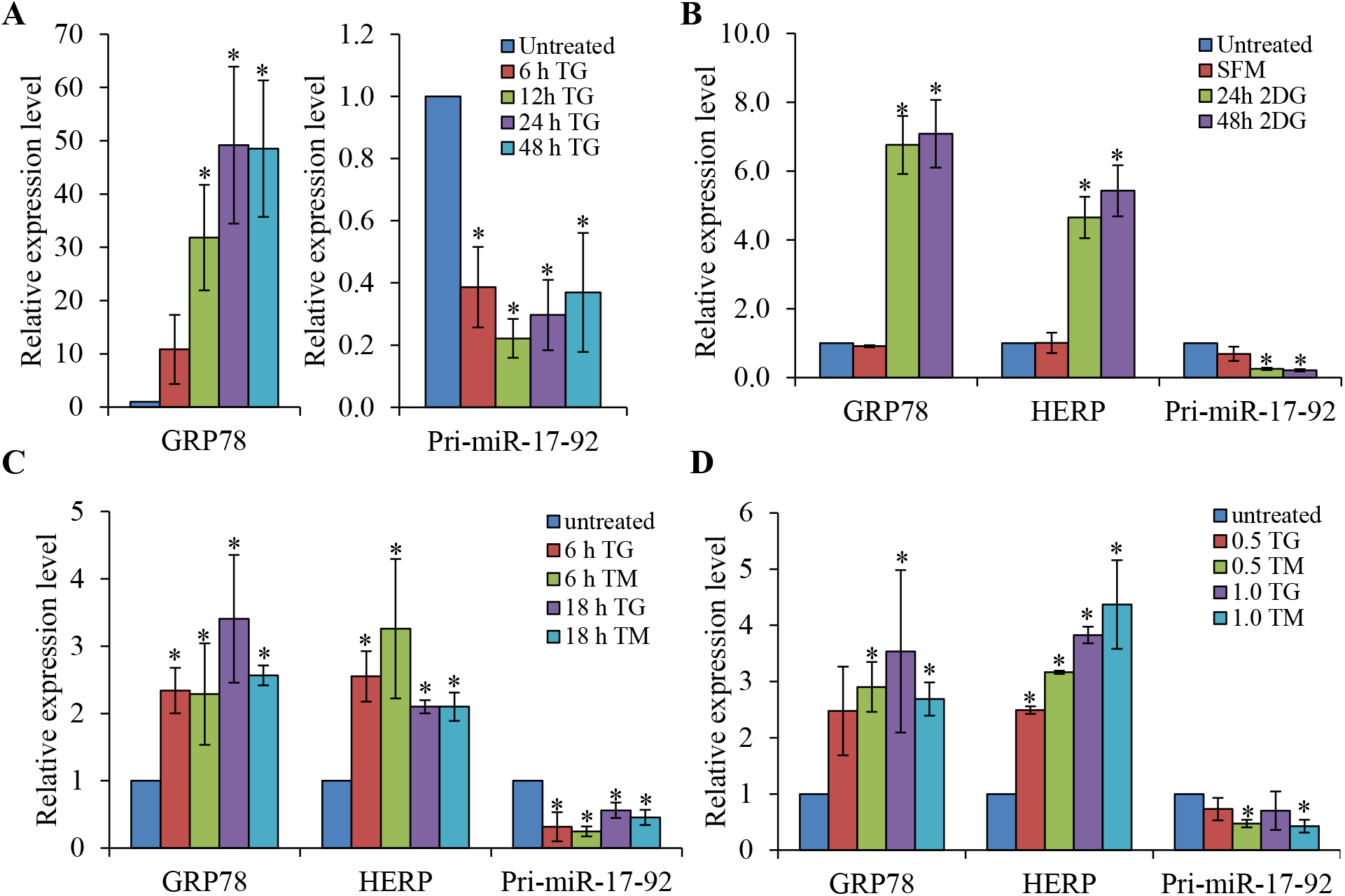
Downregulation of primary miR-17-92 transcript during conditions of ER stress. **(A)** H9c2 cells were either untreated or treated with (1.0 μM) TG for the indicated time points. The expression level of GRP78 and primary miR-17-92 transcripts was quantified by qRT-PCR, normalizing against GAPDH. Error bars represent mean ± S.D. from three independent experiments performed in triplicate. (**B**) H9c2 cells were treated with 2-deoxyglucose (1 mM) along with serum deprivation for 24 and 48 hours. The expression level of GRP78, HERP and primary miR-17-92 transcript was quantified by qRT-PCR, normalizing against GAPDH. The expression levels relative to the control are shown. Error bars represent mean ± S.D. from three independent experiments performed in triplicate. Serum free medium (SFM); 2-deoxyglucose (2DG). (**C**) MDA-MB231 cells were either untreated or treated with (1.0 μM) thapsigargin (TG) and (0.5 μg/ml) tunicamycin (TM) for the indicated time points. The expression level of GRP78, HERP and primary miR-17-92 transcript was quantified by qRT-PCR, normalizing against GAPDH. The expression levels relative to the control are shown. Error bars represent mean ± S.D. from three independent experiments performed in triplicate. (**D**) HEK-293T cells were either untreated or treated for 24 hours with (0.5 and 1.0 μM) thapsigargin (TG) and (0.5 and 1.0 μg/ml) tunicamycin (TM). The expression level of GRP78, HERP and primary miR-17-92 transcripts was quantified by qRT-PCR, normalizing against GAPDH. Error bars represent mean ± S.D. from three independent experiments performed in triplicate. (* P < 0.05, two-tailed paired t-test compared with untreated cells).

**Figure 2.**
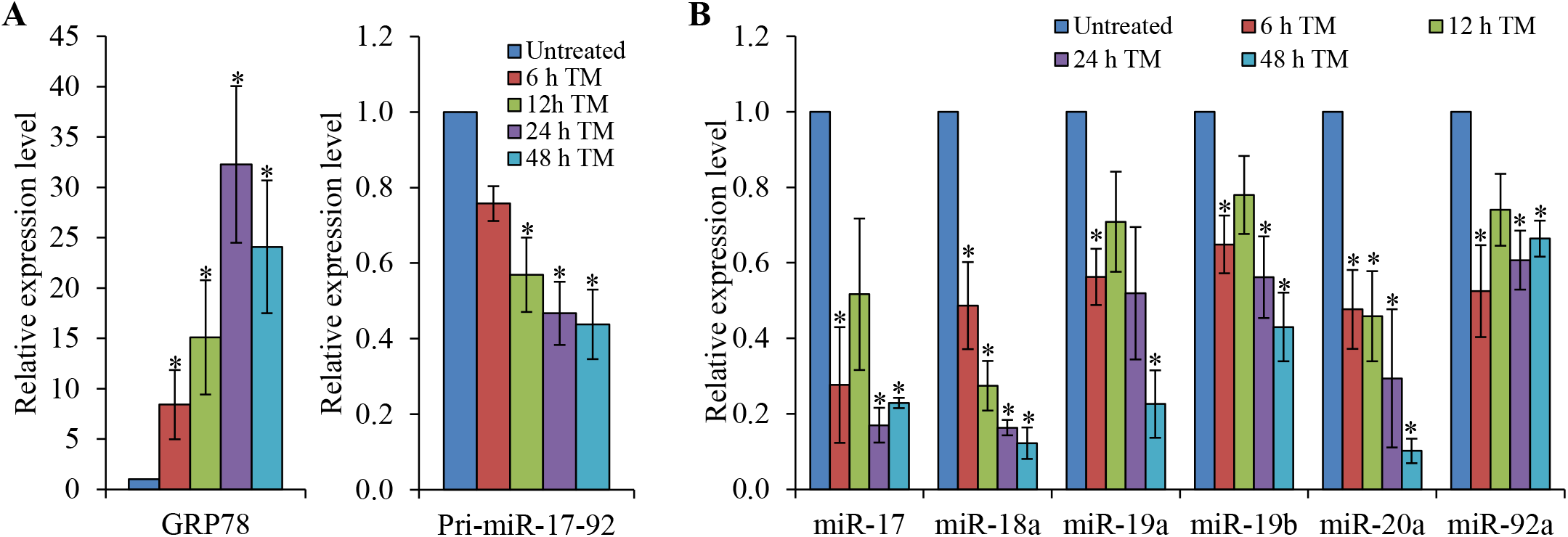
Downregulation of miRNAs belonging to miR-17-92 cluster during conditions of ER stress. Total RNA was isolated from H9c2 cells that were either untreated (Untreated) or treated with (1.0 μg/ml) tunicamycin (TM) for the indicated time points. **(A)** The expression level of GRP78 and primary miR-17-92 transcript was quantified by qRT-PCR, normalizing against GAPDH. **(B)** The expression level of member miRNAs of the miR-17-92 cluster was quantified by real-time RT-PCR, normalizing against snoRNA. Error bars represent mean ± SD from three independent experiments performed in triplicate. (* P < 0.05, two-tailed paired t-test compared with untreated cells).

### Repression of miR-17-92 promoter during ER stress is PERK-dependent

Expression of the miR-17-92 cluster has been shown to be regulated at the transcriptional level in several experimental models [14–16]. To elucidate the mechanism of downregulation of miR-17-92 cluster during conditions of ER stress we performed miR-17-92 promoter reporter assays. For this purpose we used a reporter construct having ~3700 bp DNA fragment containing the genomic locus of miR-17-92 cluster cloned at the 5’ site of the luciferase gene in the reporter pGL4.10 vector (pGL4-17-92 FL) (Figure 3A) [20]. As shown in Figure 3B, pGL4-17-92 FL reporter activity was reduced in MCF-7 cells treated with TG or TM, but no reduction in pGL4-17-92 FL reporter activity was observed upon treatment with hydrogen peroxide. The control plasmid (pGL4.10) gave much lower activity and did not show reduction in reporter activity upon treatment with TG or TM (data not shown). To explore the mechanism behind miR-17-92 cluster downregulation during ER stress, we used PERK-K618A, a dominant-negative PERK mutant that blocks transphosphorylation and activation of the PERK kinase [21, 22]. Co-expression of PERK-K618A completely inhibited the ER stress mediated downregulation of miR-17-92 cluster (Figure 3C). These data suggest that the PERK arm of the UPR is required for repression of miR-17-92 cluster during conditions of ER stress. The transcription factors ATF4, NRF2 and CHOP are activated following PERK activation during ER stress [1, 23]. Next we determined the effect of ATF4, NRF2 and CHOP co-expression on pGL4-17-92 FL reporter activity. Co-transfection of the pGL4-17-92 FL construct together with an expression vector for ATF4 and NRF2 in MCF-7 cells led to a sharp reduction in the luciferase activity, whereas co-expression of CHOP had no such effect (Figure 3D). Co-expression of ATF4, NRF2 and CHOP had no effect on the activity of control empty vector (pGL4.10) (data not shown).

**Figure 3.**
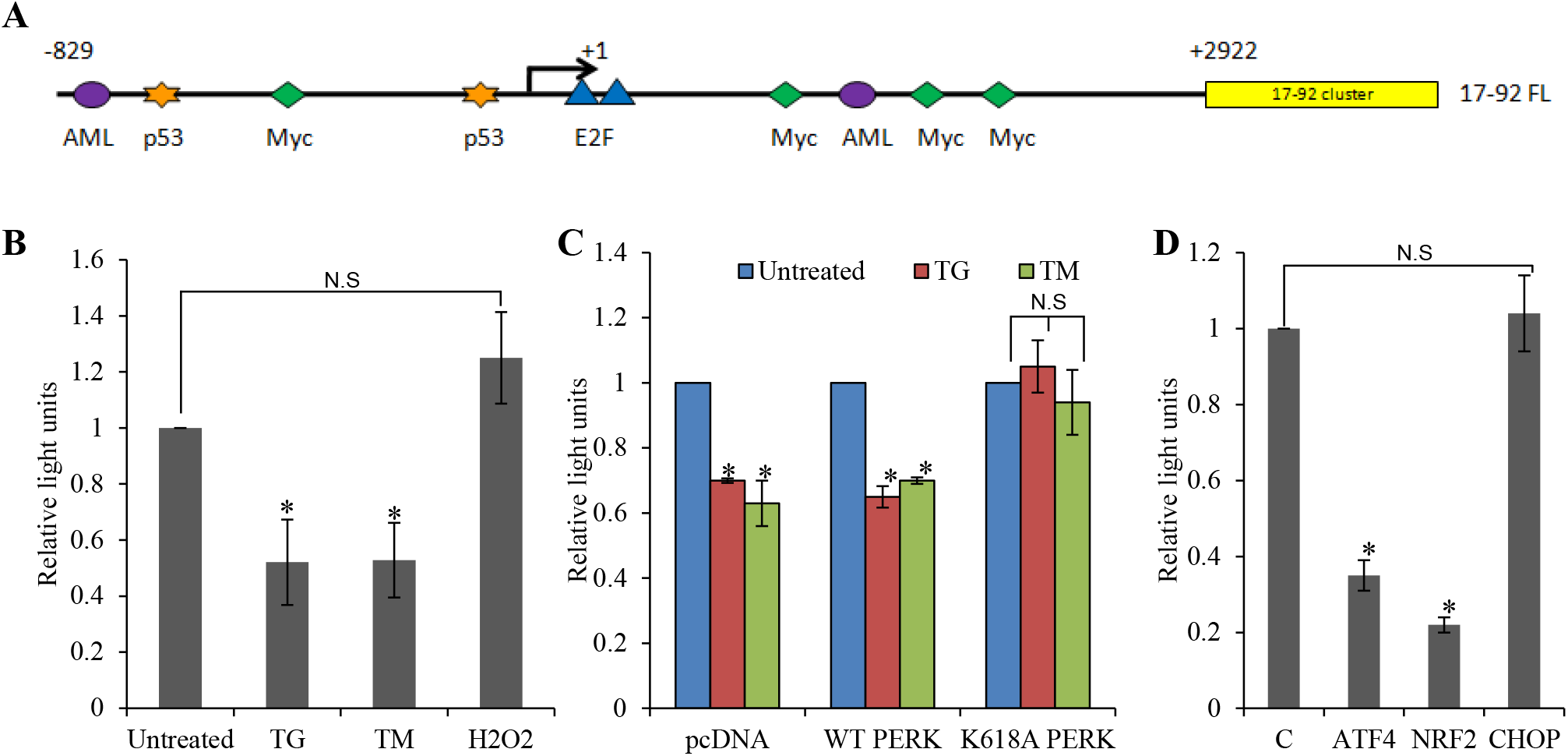
Repression of miR-17-92 promoter during ER stress is PERK dependent. **(A)** Schematic representation of the genomic region of the miR-17-92 cluster. Binding sites of the various transcription factors are shown. Arrow indicates the transcription start site. **(B)** MCF-7 cells were transfected with pGL4-17-92 FL and pRL-TK (Renilla luciferase). 24 hour post-transfection cells were left untreated or treated with (2.0 μM) thapsigargin (TG) and (2.0 μg/ml) tunicamycin (TM) or (600 μM) hydrogen peroxide (H2O2). Luciferase activity was measured 48 hour post-transfection using Dual-Glo assay system (Promega) and normalized luciferase activity (Firefly/Renilla) is shown. Error bars represent mean ± SD from three independent experiments performed in duplicate. **(C)** MCF-7 cells were transfected with pGL4-17-92 FL and pRL-TK (Renilla luciferase) in combination with pcDNA3 (pcDNA), wild-type PERK (WT PERK) or dominant-negative PERK (K618A PERK) expression plasmids. 24 hour post-transfection cells were left untreated or treated with (2.0 μM) thapsigargin (TG) and (2.0 μg/ml) tunicamycin (TM). Luciferase activity was measured 48 hour post-transfection using Dual-Glo assay system and normalized luciferase activity (Firefly/Renilla) is shown. Error bars represent mean ± SD from three independent experiments performed in duplicate. **(D)** MCF-7 cells were transfected with pGL4-17-92 FL and pRL-TK (Renilla luciferase) in combination with the control pcDNA3 (C), wild-type ATF4, NRF2 or CHOP expression plasmids. Luciferase activity was measured 24 hour post-transfection using Dual-Glo assay system and normalized luciferase activity (Firefly/Renilla) is shown. Error bars represent mean ± SD from three independent experiments performed in duplicate (* P < 0.05, two-tailed paired t-test compared with control cells).

To map the region in the miR-17-92 promoter that responds to the ER stress-mediated repression, we used promoter constructs with different lengths of the miR-17-92 5’ flanking region [-829 to +2922 (17-92FL; +1492 to + 2922 (17-92/1); +2626 to +2922 (17-92)] cloned into the promoterless luciferase pGL vector (Figure 4A) [16, 20, 24]. When transfected cells were exposed to ER stress conditions, the activities of the 17-92FL, 17-92/1 and 17-92 promoter constructs were greatly reduced in MCF-7 cells (Figure 4B). Next we determined the effect of ATF4 and NRF2 co-expression on the activity of 17-92FL, 17-92/1 and 17-92 reporter constructs. We observed significant reduction in the activity of 17-92FL, 17-92/1 and 17-92 promoter constructs upon co-expression of ATF4 or NRF2 (Figure 4C). These results suggest the presence of a cis-regulatory element responsive to ATF4 and NRF2 in the promoter region of the miR-17-92 cluster at position +2626 to +2922 (17-92) relative to the transcription start site. To elucidate the molecular mechanism of miR-17-92 repression, we transfected MCF-7 cells with GFP, ATF4 or NRF-2 expression plasmids and performed ChIP assays targeting the promoters of miR-17-92, HO-1 and CHOP. The ChIP assay confirmed the direct regulation of HO-1 (a bonafide NRF2-responsive gene) [25] by NRF2. The ChIP assay further demonstrated that NRF2 binds directly to the promoter of miR-17-92 (Figure 4D) the results for recruitment of ATF4 to miR-17-92 promoter were equivocal. ATF4 was recruited to the CHOP promoter (Figure 4D). CHOP, a known UPR-responsive gene, is confirmed here to be regulated by ATF4. Transcription factor binding analysis using Genomatix software identified a putative Nrf2-binding site (CACTTCCAGT) from +2699 to + 2708 bp in the miR-17-92 promoter [26]. Collectively, these results suggest that NRF2 can act as a transcriptional repressor for the miR-17-92 promoter. The molecular details of ATF4-mediated repression of the miR-17-92 gene remains to be elucidated. ATF4 has been described as both a negative regulator of CRE-dependent transcription [27] and a positive regulator of transcription [28]. One possibility is that ATF4 may repress the miR-17-92 promoter transcription due to squelching.

**Figure 4.**
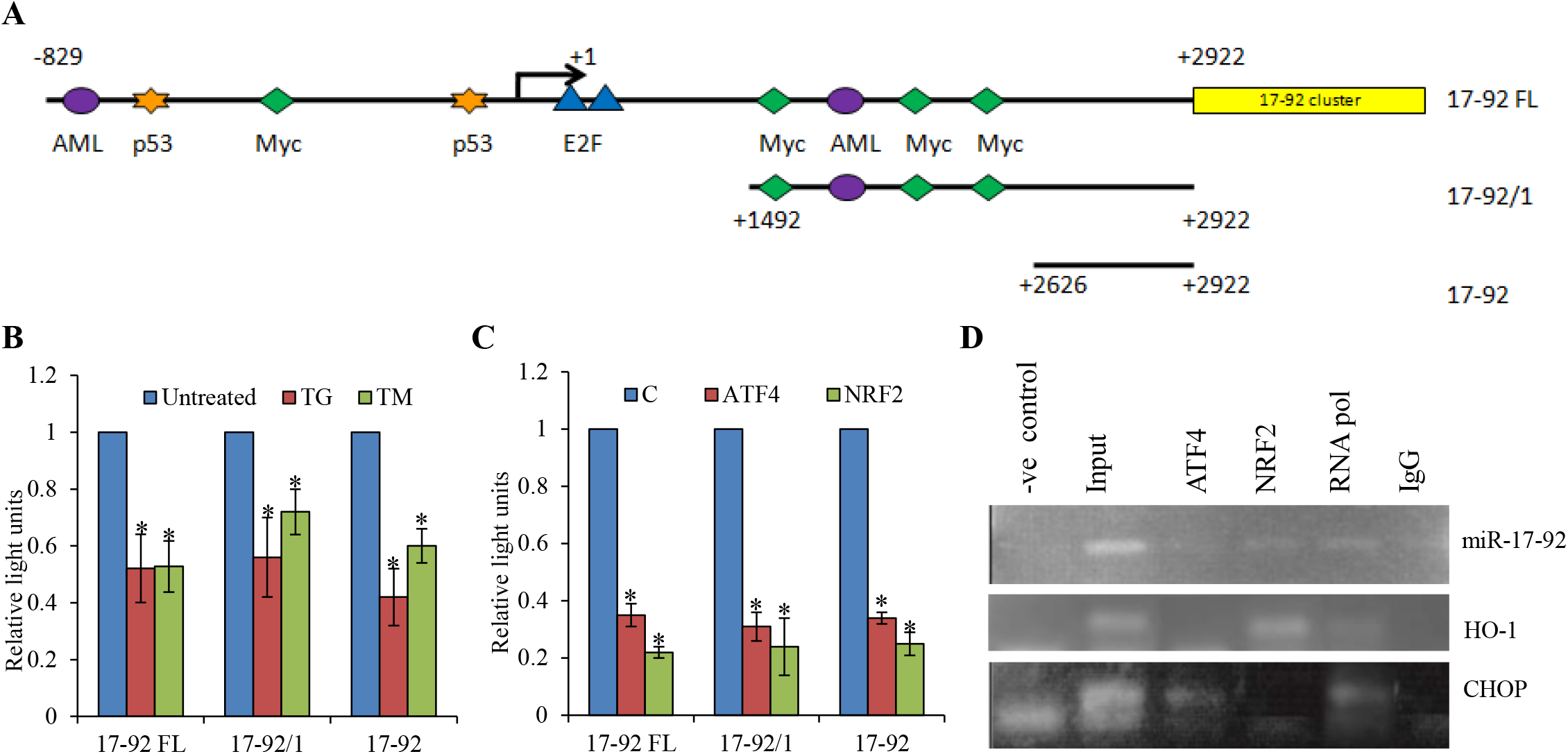
Mapping the region within miR-17-92 promoter that responds to Nrf2-mediated repression. **(A)** Schematic representation of miR17-92 cluster promoter deletion constructs is shown. **(B)** MCF-7 cells were transfected with indicated miR-17-92 promoter reporter constructs along with pRL-TK (Renilla luciferase, Promega). 24 hour post-transfection cells were either left untreated or were treated with (2.0 μM) thapsigargin (TG) and (2.0 μg/ml) tunicamycin (TM). Luciferase activity was measured 48 hour post-transfection using Dual-Glo assay system (Promega) and normalized luciferase activity (Firefly/Renilla) is shown. Error bars represent mean ± SD from three independent experiments performed in duplicate. **(C)** MCF-7 cells were transfected with indicated miR-17-92 promoter reporter constructs along with pRL-TK (Renilla luciferase, Promega) in combination with the control pcDNA3 (C), wild-type ATF4 or NRF2 expression plasmids. Luciferase activity was measured 24 hour post-transfection using Dual-Glo assay system and normalized luciferase activity (Firefly/Renilla) is shown. Error bars represent mean ± SD from three independent experiments performed in duplicate. **(D)** ChIP assays demonstrated that NRF2 bound directly on the miR-17-92 gene promoter. ChIP assay was performed with antibodies directed against either ATF4 or NRF2. Mouse IgG and RNA Pol II antibodies were used as controls for the ChIP assay. DNA was extracted from the precipitates and a DNA fragment from the miR-17-92 gene promoter sequence was amplified by PCR. DNA fragments from either HO-1 or CHOP were also amplified as controls. The PCR products were resolved on agarose gels and stained with ethidium bromide. (* P < 0.05, two-tailed paired t-test compared with control cells).

We next used the photosensitizer hypericin which accumulates prevalently in the ER membrane and upon light exposure generates reactive oxygen species (ROS), causing a loss-of function of the Ca2+-ATPase pump (SERCA) and consequent ER stress [29]. Hypericin-mediated photo-oxidative (phOx) ER damage induces PERK-dependent cell death [30]. We determined the effect of phOx-mediated ER stress on the expression level of representative miRNAs belonging to miR-17-92 cluster in wild-type and PERK-/- MEFs. We found that phOx-mediated ER stress led to a significant decrease in the levels of miR-17, miR-18a and miR-20a in wild-type but not in PERK-/- MEFs (Figure 5A). Next we determined the effect of ectopic ATF4 and NRF2 on the expression level of representative miRNAs belonging to the miR-17-92 cluster (miR-17, miR-18a and miR-19a). We found that overexpression of both ATF4 and NRF2 in PC12 cells led to a significant decrease in the levels of miR-17, miR-18a and miR-20a (Figure 5B-C). Taken together, these results show that ATF4 and NRF2 expression can repress expression of miRNAs belonging to the miR-17-92 cluster independent of ER stress.

**Figure 5.**
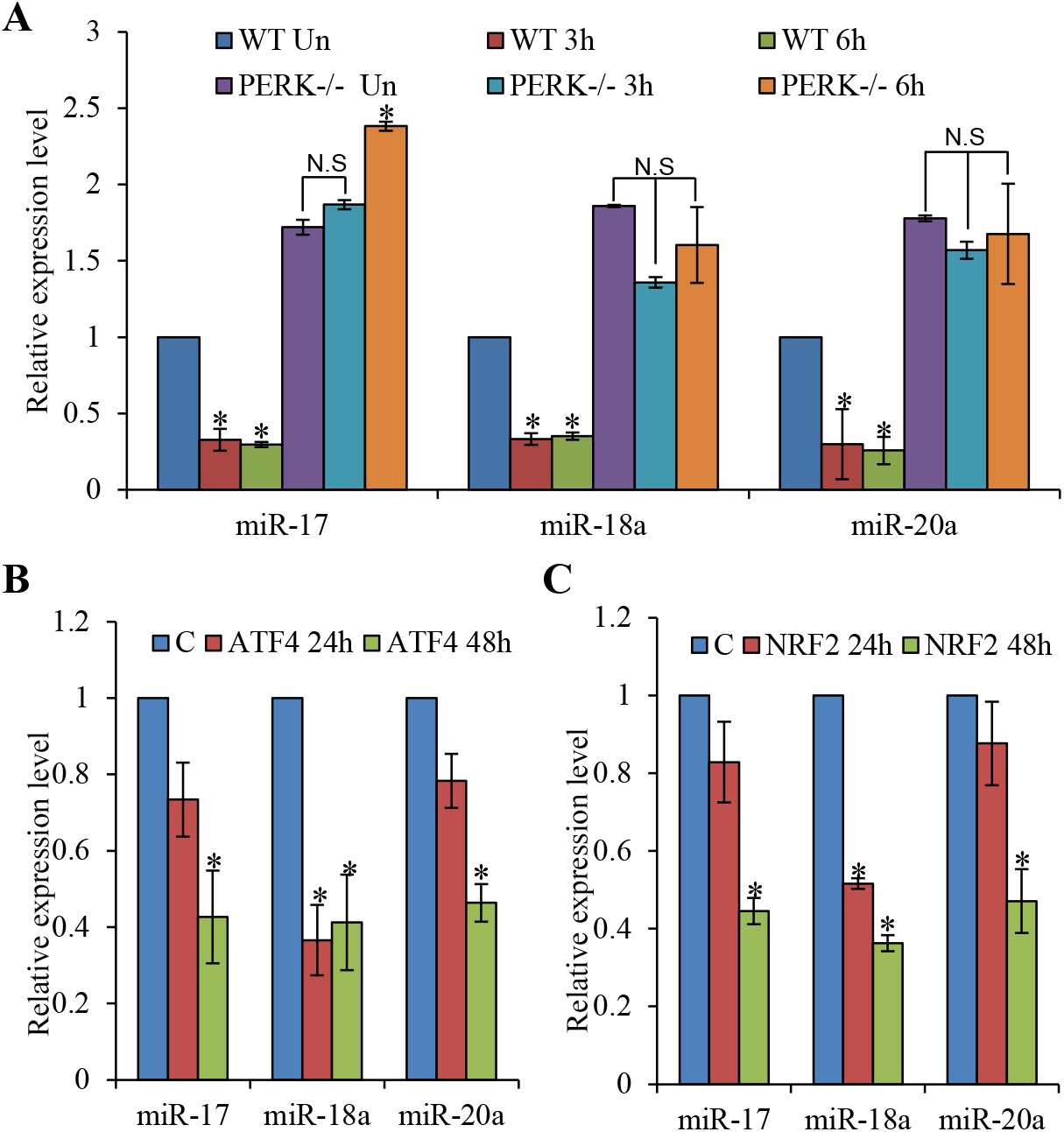
PERK-dependent regulation of miR-17-92 cluster during conditions of ER stress. (A) Wild-type (WT) and PERK knockout (PERK-/-) MEFs were exposed to phOx stress (200nM hypericin for 2h, 2,7J/cm2). Total RNA was isolated and expression level of miR-17, miR-18a and miR-20a was quantified by real-time RT-PCR, normalizing against snoRNA. Error bars represent mean ± SD from two independent experiments performed in triplicate. (B) PC12 cells were transfected with the control pcDNA3 (C), wild-type ATF4 or NRF2 expression plasmids and total RNA was isolated at the indicated time points. Expression levels of miR-17, miR-18a and miR-20a were quantified by real-time RT-PCR, normalizing against snoRNA. Error bars represent mean ± SD from three independent experiments performed in triplicate. (* P< 0.05, Two-tailed paired t-test compared to control samples).

### MiR-17-92 cluster protects against UPR-induced cell death

To determine the role of miR-17-92 cluster on ER stress-induced apoptosis, we evaluated the effect of miR-17-92 cluster overexpression on ER stress-induced apoptosis. Next we generated clones of SH-SY5Y and H9c2 cells expressing miR-17-92 to evaluate its role in ER stress-induced apoptosis. For this purpose SH-SY5Y and H9c2 cells were transduced with lentivirus engineered to produce RFP and miR-17-92. The control and miR-17-92 overexpressing clones of SH-SY5Y (Figure 6A) and H9c2 (Figure 6E) showed expression of RFP. The miR-17-92 expressing clones of SH-SY5Y (Figure 6B) and H9c2 (Figure 6F) cells showed increased expression of miRNAs belonging to miR-17-92 cluster as compared to the control clones. However, we observed variations the level of individual miRNAs in miR-17-92 expressing clones of SH-SY5Y (Figure 6B) and H9c2 (Figure 6F) cells suggesting cell-type dependent processing of primary transcript. Next, we evaluated whether expression of miR-17-92 cluster can affect the sensitivity to ER stress-induced cell death. We found that ER stress-induced apoptosis was attenuated in miR-17-92 expressing clones of SH-SY5Y (Figure 6C) and H9c2 (Figure 6G) cells as compared with control clones. Western blot analysis revealed that treatment with TG and TM induced processing of caspase-3 in control and miR-17-92 overexpressing clones of SH-SY5Y (Figure 6D) and H9c2 (Figure 6H) cells. Notably there was decreased processing of caspase-3 in miR-17-92 expressing cells as compared to control cells (Figure 6D & H). Thus, overexpression of miR-17-92 cluster provides resistance against ER stress-induced apoptosis.

**Figure 6.**
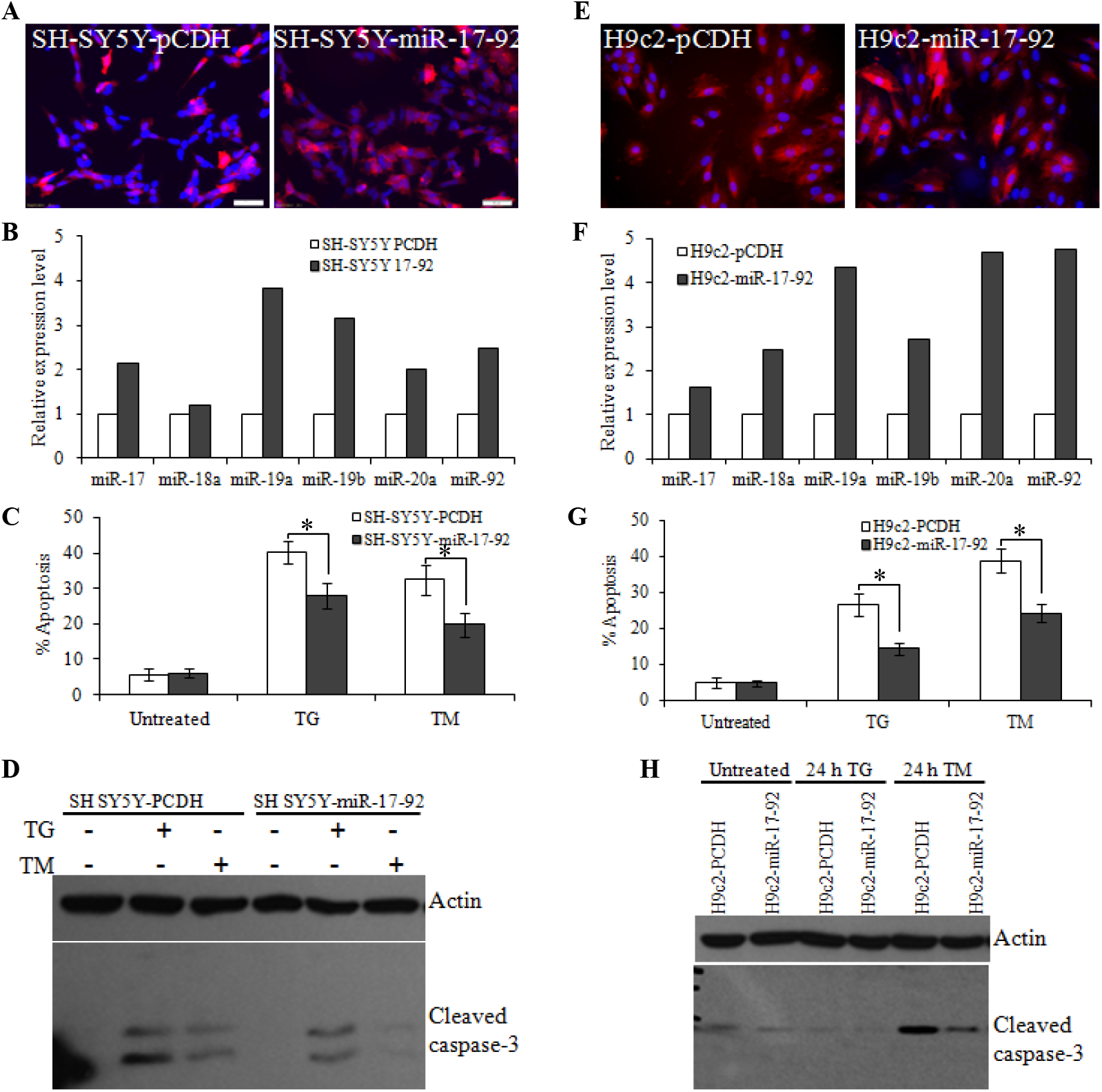
Effect of miR-17-92 cluster on UPR-mediated cell death. (A) SH-SY5Y-PCDH and SH-SY5Y-miR-17-92 cells were grown on cover slips for 24 h and expression of RFP was monitored after fixing and mounting the cells in mountant containing DAPI. (B) Total RNA was isolated from SY5Y-PCDH and SH-SY5Y-miR-17-92 cells and expression level of miRNAs belonging to miR-17-92 cluster was quantified by qRT-PCR, normalizing against snoRNA. (C) SH-SY5Y-PCDH and SH-SY5Y-miR-17-92 cells were treated with (0.5 μM) TG and (0.5 μg/ml) TM for 24 h. After treatment the apoptotic nuclei were determined as described in Materials and Methods. Error bars represent mean ± S.E. from three independent experiments performed in triplicate. (D) SH-SY5Y-PCDH and SH-SY5Y-miR-17-92 cells were treated with (1.0 μM) TG and (1.0 μg/ml) TM for 24 h. Western blotting of total protein was performed using antibodies against cleaved caspase-3 and actin. (E) H9c2-PCDH and H9c2-miR-17-92 cells were grown on cover slips for 24 hours and expression of RFP was monitored after fixing and mounting the cells in mountant containing DAPI. (F) Total RNA was isolated from H9c2-PCDH and H9c2-miR-17-92 cells and expression level of miRNAs belonging to miR-17-92 cluster was quantified by qRT-PCR, normalizing against snoRNA. (G) H9c2-PCDH and H9c2-miR-17-92 cells were treated with (1.0 μM) TG and (1.0 μg/ml) TM for 24 h. After treatment the apoptotic nuclei were determined as described in Materials and Methods. Error bars represent mean ± S.E. from three independent experiments performed in triplicate. (H) H9c2-PCDH and H9c2-miR-17-92 cells were treated with (1.0 μM) TG and (1.0 μg/ml) TM for 24 h. Western blotting of total protein was performed using antibodies against cleaved caspase-3 and actin. (* P < 0.05, two-tailed paired t-test compared with control cells).

Our study indicates that miRNAs belonging to the miR-17-92 cluster have a pivotal role in the UPR. The expression of these miRNAs is markedly downregulated during ER stress conditions. The repression of miR-17-92 cluster during ER stress is dependent on PERK signalling. Furthermore, our results show that miR-17-92 cluster is a novel target for ATF4 and NRF2-mediated transcriptional repression under conditions of ER stress. Ectopic expression of ATF4 or NRF2 can lead to a decrease in levels of miRNAs comprising the miR-17-92 cluster as well as a reduction in miR-17-92 promoter reporter activity. Their functional role during ER stress was demonstrated by overexpression experiments, whereby an increased expression of miR-17-92 cluster inhibits ER stress-induced apoptosis.

The miR-17-92 cluster is located on chromosome 13 in the locus of the non-protein-coding gene MIR17HG (the miR-17/92 cluster host gene). Ancient gene duplications have given rise to two miR-17-92 cluster paralogs in mammals: the miR-106b-25 and the miR-106a-363 cluster, respectively [13]. Together these three miRNA clusters represent a combined total of 15 miRNAs that form four ‘seed’ families: the miR-17 family, the miR-18 family, the miR-19 family and the miR-92 family. Furthermore the miR-17-92 and miR-106b-25 clusters have been shown to be transcriptionally regulated by the MYC and E2F family of transcription factors [13]. We have recently shown that the miR-106b-25 cluster is downregulated during ER stress, in a PERK-dependent manner, and contributes to optimum induction of BIM and ER stress-induced cell death [12]. Thus both miR-17-92 and miR-106b-25 clusters are regulated by similar mechanisms during UPR and can modulate the resistance to ER stress-induced cell death.

What is the biological significance of transcriptional repression of the miR-17-92 cluster by ATF4 and NRF2? Transcription factors, ATF4 and NRF2 are activated during ER stress in a PERK-dependent manner. Genetic and pharmacological experiments have demonstrated that PERK signalling can confer both protective and proapoptotic effects in response to ER stress [4, 31, 32]. For instance, genetic deletion of PERK sensitizes the cells to ER stress-induced apoptosis [31]. However, sustained PERK signalling has been shown to impair cell proliferation and promote apoptosis [33]. ATF4 has been shown to act as a pro-death transcriptional regulator in the nervous system that propagates death responses to oxidative stress *in vitro* and to stroke *in vivo* [34]. ATF4 also mediates ER stress-induced cell death of neuroectodermal tumour cells in response to fenretinide or bortezomib [35]. Further sustained activation of NRF2 in ATG5-deficient mouse livers due to p62 mediated stabilization of NRF2 has been reported to be a major cause of toxicity in autophagy-impaired livers [36]. However, the molecular mechanisms by which ATF4 and NRF2 exert their pro-apoptotic effects are not clearly understood. The data presented here provides the molecular mechanism underlying the PERK-mediated induction of cell death.

## Acknowledgements

We are grateful to the Technical Officers and administrative team in Pathology, School of Medicine, NUI, Galway. We would like to thank Maria Ryan and Ayswaria Deepti for technical assistance. This publication has emanated from research conducted with the financial support of Health Research Board (grant number HRA_HSR/2010/24) to S.G.

## Authors’ contributions

DR and AG performed experiments, analysed the data and wrote the manuscript. DR performed experiments, analysed the data and wrote the manuscript. KC performed experiments, analysed the data. LF provided technical and material support. PA provided the RNA samples from PERK+/+ and PERK-/- MEFs after hypericin-mediated photo-oxidative (phOx) ER damage. AS analysed the data and supervised the work. SG conceived the project, performed experiments, analysed the data, supervised the work and wrote the manuscript. All authors read and approved the final manuscript.

